# Enzyme Immunoassays May Be Inaccurate: Familiar Correlates of Male Testosterone (T) Replicate in Britons when T is Measured by Mass Spectrometry but Do Not Replicate in Americans When T is Measured by Enzyme Immunoassay

**DOI:** 10.1101/351734

**Authors:** Allan Mazur, Soazig Clifton

## Abstract

Several correlations have been reported between men’s testosterone (T) and other characteristics, e.g., T declines during the day, declines with obesity, and declines with advancing age. We asked if these relationships hold in older men when T is assayed from saliva. Here seven familiar correlations are tested among older American men, their salivary T measured by enzyme immunoassay (EIA). Some correlations can also be tested among older British men, their salivary T assayed by liquid chromatography-tandem mass spectroscopy (LC-MS/MS), a technique noted for its specificity. Most of our attempts at replication failed in the American data but succeeded in the British data. We conclude that failure to replicate in Americans is likely due to inaccuracy of EIA values for T, and that most T correlations hold true in older men when salivary T is accurately measured by LC-MS/MS.

## INTRODUCTION

Well documented correlates of men’s testosterone (T) include its mean level peaking in the teens to twenties and thereafter declining with age; a decrease in T with obesity; and diurnal decline in T from a high level in the morning. But these (and other) relationships, typically observed among samples including young and middle-aged men, may not hold among older men. “Endocrine function undergoes major changes during aging... Specifically, alterations in hormonal networks and concomitant hormonal deficits/excess, augmented by poor sensitivity of tissues to their action, take place. As the hypothalamic–pituitary unit is the central regulator of crucial body functions, these alterations can be translated in significant clinical sequelae” (Diamanti-Kandarakis et al. 2017).

Here we ask if seven previously-reported correlates of T, usually measured from serum, are still seen in older men when T is assayed from saliva. We tested these correlations in two large datasets, one American and one British. American data were collected by the National Social Life, Health & Aging Project (NSHAP), a two-wave panel (years 2006 and 2011) of older Americans ranging in age from 57 to 85 at Wave 1. T was measured in each wave by enzyme immunoassay (EIA) of saliva samples collected at the time of interview.

British data were collected in a single wave of the National Survey of Sexual Attitudes and Lifestyles: 2010-2012 (Natsal-3), which assayed salivary T by liquid chromatography-tandem mass spectroscopy (LC-MS/MS), a technique noted for its specificity. Natsal-3 did not include all the variables used from NSHAP so not as many correlations could be tested.

Several validation studies show salivary T to be correlated with “free” T in serum, i.e., with that small portion of total T not bound to protein and thus physiologically active (Shirtcliff et al. 2002; Keevil et al. 2014; Fiers et al. 2014). Biobehavioral researchers increasingly collect saliva rather than blood since saliva is cheaper and easier. Among several assay techniques, EIA has become a workhorse of behavioral studies because it is quick, convenient, inexpensive, and does not require radioactive material. Recently, however, Welker et al. (2016) presented results that question the accuracy of EIA for measuring salivary T.

### PREVIOUSLY REPORTED CORRELATES OF MALE T

Without presenting a full literature review, we note correlations previously reported, which can be tested on one or both of our datasets. For example, it is now well established that in men, T has a diurnal rhythm, that low T is associated with obesity, and that T usually declines as men age. These head the list of propositions tested here in older men, which also contains other T correlations reported less frequently or consistently:

1. <u>Diurnal</u>: Male T has a diurnal rhythm, high in the morning, lower as the day goes on (Dabbs 1990; Wittert 2014; Diver et al. 2003; Luboshitzky et al. 2003).
2. <u>Obesity</u>: Low T is associated with obesity (e.g., Vermeulon et al. 1999; Derby et al. 2006; Wu et al. 2008; Kim et al., 2012; Camacho et al. 2013; Shi et al. 2013; Bann et al. 2015; Saad et al. 2016; Clifton et al. 2016).
3. <u>Age</u>: At least in industrial societies, men past the age of thirty tend to show declining T (Vermeulon et al. 1999; Travison et al. 2006; Andersson et al. 2007; Liu et al. 2007; Wu et al. 2008; Mazur 2009; Bann et al. 2015; Clifton et al. 2016).
4. <u>Consistency across waves</u>: Basal T is consistent from year to year, despite fluctuations during the day. For example, among Australian men, r = .73 for serum T measured five years apart (Shi et al. 2013, Supplemental Figure 1).
5. <u>Sexuality</u>: Lower T is associated with lower sexuality, variously measured (Yeap et al. 2012; Corona et al. 2017; Rizk et al. 2017; Rosen et al. 2017). With the rising popularity of testosterone replacement therapy (TRT), there has been much attention to associations between T level and male mortality or morbidity. Some observers doubt TRT’s efficacy as a rejuvenator and others warn of health risks (e.g., Handelsman, 2017; Finkel et al. 2014; Vigen et al. 2013). Most prominent has been the issue of whether high T improves longevity or shortens it by promoting cardiovascular risks (Araujo et al. 2011; Shores et al. 2012; Yeap et al. 2012; Vigen et al. 2013; Pastuszak et al. 2017). Therefore, two additions propositions for test are:
6. <u>Diabetes</u>: High T is associated with low HbA1c and low incidence of Type 2 diabetes in men (Selvin et al. 2007; Haider et al. 2014; Yassin et al. 2014), though finer grain analysis casts some doubt on the validity of earlier reports (Mazur et al. 2013).
7. <u>Mortality and morbidity</u>: Knowing which men in Wave 1 of NSHAP died before Wave 2 provides an opportunity to test if T in Wave 1 predicted death. Several measures of health in both waves, apart from diabetes, allow tests for their relationships to T.

## METHODS

### Two Surveys: American NSHAP and British Natsal-3

The Institutional Review Board of Syracuse University has determined that this research meets the organization’s ethical standards and qualifies for exemption from human subject regulation.

The National Social Life, Health, and Aging Project (NSHAP) is a longitudinal, population-based study of health and social factors, aiming to understand the well-being of older, community-dwelling Americans by examining the interactions among physical health and illness, medication use, cognitive function, emotional health, sensory function, health behaviors, social connectedness, sexuality, and relationship quality (https://www.nia.nih.gov/research/resource/national-social-life-health-and-aging-project-nshap). Wave 1 (in 2005) included over 3,000 interviews of men and women drawn to be nationally representative of older Americans born between 1920 and 1947 (aged 57 to 85 at Wave 1). Wave 2 (in 2010) had nearly 3,400 interviews including Wave 1 respondents, plus people drawn for Wave 1 who were not interviewed at that time, plus spouses or cohabiting romantic partners.

The National Survey of Sexual Attitudes and Lifestyles (Natsal-3) is a multi-wave probability-sample study of British men and women to investigate associations among demographic characteristics, lifestyle, general health, and reported health conditions (Erens et al. 2014). Only the 2010-12 wave is relevant here because it was the first to collect saliva and assay for T. This wave included 15,162 men and women, ages 16-74 years. Thus, NSHAP has two waves with T measured in older Americans, while Natsal-3 has one wave with T measured in a wide age range of adult Britons.

The present analysis is limited to male respondents because most propositions listed above apply to or are mostly reported for men, and because men’s higher levels of T are easier to measure accurately. To nearly equate age ranges in the US and UK studies, analysis is limited to British respondents age 57 years or older. Cultural US-UK differences in the early 21^st^ century are ignored. For present purposes, the critical distinction between studies is their different methods of assaying salivary T: EIA in the American sample, LC-MS/MS in the British sample.

Saliva samples for NSHAP were collected at the time of interview, which was variable through the day. Respondents were asked to provide saliva by passive drool into a code-labeled polypropylene vial through a household plastic straw, following procedures recommended by Salimetrics, LLC (Carlsbad, CA 92008). Wave 1 specimens were transported from the interview to a freezer using cold packs, then stored in a freezer until shipped on dry ice to Salimetrics, which upon receipt stored them at −80°C. On the day of assay, specimens were thawed completely, vortexed, and centrifuged. Clear samples were pipetted into wells. Salimetrics kits were used for enzyme immunoassay. The assay range was 1.0 pg/ml to 600 pg/ml. Intra-assay precision was determined from the mean of 8 replicates at high and low T levels. The average intra-assay coefficient of variation was 3.3% and 6.7% for high and low levels. Inter-assay precision was determined from the mean of averaged duplicates for 10 separate runs at high and low testosterone levels. The average inter-assay coefficient of variation was 5.1% for high and 9.6% for low testosterone levels. Saliva samples were run in duplicate. Results for each subject were acceptable when the coefficient of variation (%CV) between the duplicates was <15%. In instances where the %CV between duplicates was >15%, results were accepted if the absolute difference between result 1 and result 2 was <8 pg/ml. If these criteria were not met, saliva was assayed again. Values greater than the upper assay limit of 600 pg/ml were run on dilution to bring the readings within the accepted range. Procedures for Wave 2 were similar, including the use of Salimetrics EIA kits, however these samples were shipped to Germany for assay by C. Kirschbaum at the University of Dusseldorf (Mendoza et al. 2007; Gavrilova and Lindau 2009; O’Doherty et al. 2014; Kozloski et al. 2014).

For men, the correlation between the first two (i.e., duplicate) T values is r = 0.99 in Wave 1 and r = 0.97 in Wave 2 (both p <.001). Duplicates are averaged to establish each respondent’s mean T except when a duplicate is not recorded, in which case the one recorded value is taken as mean T. In Wave 1, six men are excluded because of implausibly high mean T (> 1000 pg/ml).

The British Natsal-3 obtained a usable saliva sample from 4,128 respondents (1675 men, 2453 women), who were instructed to provide a saliva sample before 10:00 on the morning after their interview. They were asked not to brush their teeth, eat or chew before giving the sample, then drool into a polystyrene vial, and post the sample on the day of collection to the Department of Clinical Biochemistry, Glasgow Royal Infirmary (GRI), where they were prepared and frozen at −80°C until analysis. Assays were performed at the Biochemistry Department at University Hospital South Manchester, using a newly developed and validated liquid chromatography tandem mass spectrometry technique (MacDonald et al. 2011).

NSHAP reports T levels in metric units, Natsal-3 in molar units. The conversion is 1 pmol/l = 0.3 pg/ml (https://www.menshormonalhealth.com/hormone-unit-conversion-calculator.html).

### Variables

Beside T, the following variables available in NSHAP are listed in order of the propositions to be tested. Not all these variables were available from Natsal-3, either because they were not collected or, for sexuality, were collected but not released to us. We accepted variables in the public datasets at face value, having no control over their collection.

1. <u>Diurnal</u>: Time of day (24-hour clock) when saliva was collected is used to check diurnal rhythm of T.
2. <u>Obesity</u>: Three measures of obesity are used: body mass index (BMI), waist circumference, and interviewer’s rating of respondent’s body shape on a 4-point scale from thin to obese.
3. <u>Age</u>: Age at each wave is given in years.
4. <u>Consistency across waves</u>: T values for men in NSHAP who provided usable saliva on both waves were correlated to assess consistency across waves.
5. <u>Sexuality</u>: NSHAP contains several variables on sexuality, from which a subset was chosen with relatively high response rate and variability: How important is sex in your life? How often do you think about sex? How often do you masturbate? During the last 12 months, was there a period of several months…when you lacked interest in having sex? ...when you were unable to climax? ...when you had trouble getting or maintaining an erection? Married men only: In the past 12 months, how often did you have sex?
6. <u>Diabetes</u>: HbA1c and self-report of diabetes are recorded in both the American and British studies.
7. <u>Mortality and morbidity</u>: NSHAP records those respondents in Wave 1 who died before Wave 2. These may be compared to survivors, controlling on other risk factors. (Of 1,454 men with T measured in the first wave, 231 died before Wave 2.) Also, each wave contains several measures of morbidity, in addition to HbA1c and diabetes, from which these offered good response, variability, and pertinence: Would you say your health is excellent, very good, good, fair, or poor? Has a medical doctor ever told you that you had a heart attack? Have you ever been treated for heart failure? Has a medical doctor every told you that you have high blood pressure?

### Statistics

Most results reported here for NSHAP come from our regression models, including logistic regression when the dependent variable is dichotomous, with control variables where appropriate. Scatterplots are employed for some visual displays. Results are either cross-sectional, for each wave, or longitudinal. For brevity, we report relevant summary statistics, focusing especially on statistical significance, or lack of it.

Most propositions were first tested on 841 men who gave usable T in both waves. This produced unexpectedly few significant relationships by the common criterion of p = .05. As a cautionary step, propositions were tested again with samples enlarged by also including those men who gave usable T only on one wave. This increased sample sizes to 1,454 men for Wave 1, and 1,369 men for Wave 2. Of course, relationships were more likely to reach significance with these larger samples. It is worth emphasizing that correlations reaching significance with so many respondents may be slight in magnitude, having little substantive importance.

The distribution of T values is slightly skewed in Wave 1 (skew = 1.4) and considerably skewed in Wave 2 (skew = 6.3). A transformation to lnT corrected this for Wave 2 (skew = −0.1) while having little effect on Wave 1. Analyses were run with both T and lnT, usually with similar results. Occasionally a correlation reached significance with one but not the other, in which case it was treated as significant. The reason for these conservative judgements is to avoid dismissing previously-reported propositions as insignificant in NSHAP’s older men.

Statistical analyses for Britain’s Natsal-3, in which salivary T was assayed by LC-MS/MS, were carried out using STATA (version 13.1) accounting for the complex survey design (stratification, clustering, and weighting of the sample). Two weights were applied: a survey weight correcting for unequal probability of selection and differential response (by age, sex, and region) to the survey itself, and an additional saliva weight correcting for unequal probability of selection and differential response to the saliva sample (Clifton et al 2016). Multivariable linear regression was used to assess associations with mean salivary T.

Some Natsal-3 results cited here were previously reported (Clifton et al. 2016), others come from our separate analysis of men aged 57 or older. Values for salivary T were censored for very high levels so that, for each 10-year age group stratified by sex, values above the 99th percentile were assigned a value equal to that of the 99th percentile. T data for men were normally distributed.

## RESULTS

Table 1 compares basic characteristics for NSHAP (both waves) and Natsal-3 (all ages, and age 57+ years). Of note is NSHAP’s relatively low sex ratio of T, one of the issues raised by critics of EIA assays (e.g., Welker et al. 2016).

**Table 1.** Characteristics of Respondents with Usable Salivary Testosterone Values in NSHAP (both waves) and Natsal-3 (all ages and 57+ years).

|  | NSHAP Wave 1<br>(n = 1,414) | NSHAP Wave 2<br>(n = 1,369) | Natsal-3<br>(n = 1,886) | Natsal-3, Age 57+<br>(n = 564) |
| --- | --- | --- | --- | --- |
| Age range<br>(years) | 57-85 | 51-93 | 16-74 | 57-74 |
| Mean T of men<br>(pg/ml) | 85.8 (sd=46.0)<br>(n=1454) | 80.6<br>(n=1369) | 66.9<br>(n = 1599) | 48.6<br>(n =571) |
| Sex ratio of T | 1.82 | 1.80 | 6.0 | 5.9 |

Results are first reported for the American NSHAP data, with T assayed by EIA.

1. <u>Diurnal</u>: Figure 1 shows, for each wave, a scatterplot of T as a function of time of day when saliva was collected. The figure is based on 841 men with usable T on both waves. Visual inspection fails to show the expected diurnal variation, however the correlation between lnT and time of sampling is significant if small in Wave 1: r = −0.21 (p < .001). Using the enlarged sample (including men with T in only a single wave) produced consistently significant but very small correlations: r = −0.10 (p <.001) in Wave 1; r = −0.06 (p = .03) in Wave 2.

**Figure 1.**
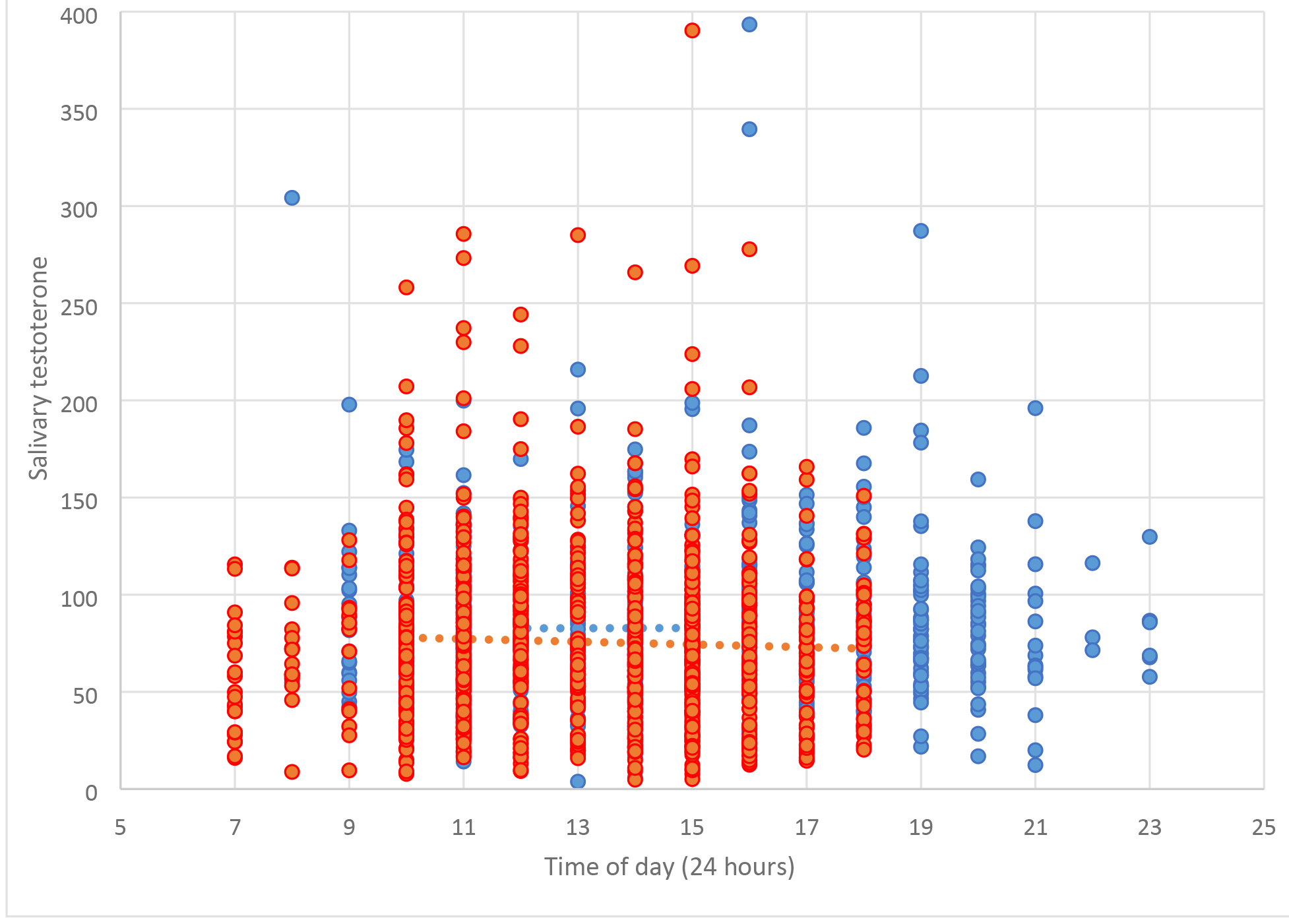
Salivary testosterone by time of day when saliva was collected, for American men in NSHAP. Values for Wave 1 are shown in red, and for Wave 2 in blue. Linear regression lines are shown for each wave.
2. <u>Obesity</u>: Whether T declines with obesity was tested on three measures: BMI, waist circumference, and interview’s evaluation of the respondent’s body shape as thin to obese. Nearly all correlations were insignificant, except that lnT is significantly if weakly correlated (inversely) to waist circumference, using the enlarged sample for Wave 1 (r = −0.06, p = .04).
3. <u>Age</u>: Figure 2 shows, for each wave, a scatterplot of T as a function of age, for 841 men with T in both waves. Visually, there is no apparent decline with age in Wave 1 and a slight decline in Wave 2. This wave difference is corroborated by calculated correlations of T and age: insignificant in Wave 1; r = −0.20 (p < .001) in Wave 2. Using the enlarged samples, correlations are significant if weak in both waves: r = −0.08 (p = .01) for Wave 1; r = −0.15 (p < .001) for Wave 2.

**Figure 2.**
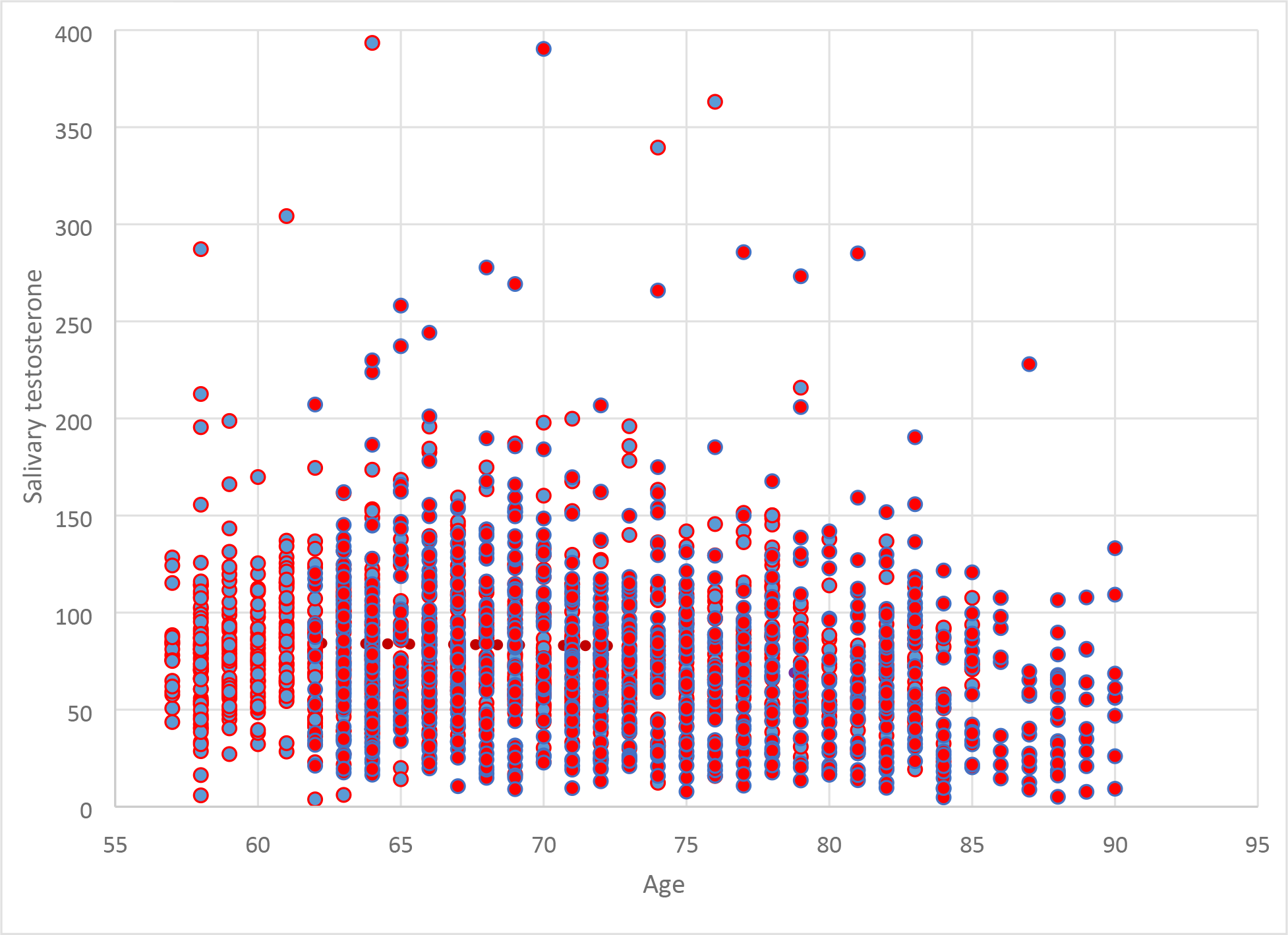
Salivary testosterone by age, for American men in NSHAP. Values for Wave 1 are shown in red, and for Wave 2 in blue. Linear regression lines are shown for each wave.
4. <u>Consistency across waves</u>: Among the 841 men with T values measured in both waves, five years apart, the correlation between waves is r = 0.23 (p < .001) for lnT; and slightly weaker for untransformed T: r =0.17 (p < .001).
5. <u>Sexuality</u>: Using enlarged samples, correlations in Wave 1 between T or lnT and all seven indicators of sexuality are insignificant, with or without controlling on age. In Wave 2, zero-order correlations are significant if small for importance of sex (r = 0.16, p < .001), thinks about sex (r = .10, p = .003), frequency of masturbation (r = .10, p < .001), and difficulty getting an erection (p = .01). Controlling on age lessens the strength and significance of these relationships. The remaining indicators of sexuality (frequency of sex, lack of interest in sex, difficulty reaching climax) are never significantly related to T. In sum, Wave 1 shows no relation between T and sexuality. Wave 2 shows significant but slight correlations with T and four of seven indicators of sexuality, which are diminished after controlling on age.
6. <u>Diabetes</u>: Even for the enlarged samples, correlations between T or lnT, on the one hand, and HbA1c or type 2 diabetes, on the other, do not approach significance in either wave. Controlling on age gives no improvement.
7. <u>Mortality and morbidity</u>: The relationship between T in Wave 1, and living or dying by Wave 2, was explored by controlling on several known risk factors: age, BMI, diagnosed diabetes, diagnosed hypertension, cigarette smoking, respondent’s self-rating of his physical health, and if respondent was aware of having heart failure. Neither T nor lnT approached significance in any of these logistic models. Unsurprisingly, when combined into a single model (n = 1139), those factors most significantly related to death were age, heart failure, self-rating of physical health (all p ≤ .001), and smoking (p = .02). BMI and diagnosis of either diabetes or hypertension approached significance (p < .1).

Overall, tests with the American NSHAP data, in which T was measured by EIA, produced weak or null findings, generally failing to replicate correlations previously reported in the literature.

Turning now to older British men in Natsal-3, for whom salivary T was assayed by LC-MS/MS, we note first that not all propositions can be tested for lack of suitable variables so results are necessarily abbreviated.

1. <u>Diurnal</u>: No opportunity to test.
2. <u>Obesity</u>: T was inversely related to measures of obesity in men of all ages, including those age 55 −74 years, though the difference between obese and normal men lessened with increasing age. In the age range 55-64, mean T of obese men (BMI>30 kg/m^2^) is 485 pg/ml, while mean T of normal weight men (BMI 18.5-25 kg/m^2^) is 635 pg/ml. In the age range 65-74, mean T of obese men is 450 pg/ml, while mean T of normal weight men is 515 pg/ml (Clifton et al. 2016: Figure 1A).
3. <u>Age</u>: British men aged 55-64 years had mean T of 558 pg/ml, while those aged 65-74 had mean T = 507 pg/ml. This decadal decline of 51 pg/ml is significant (p < .001), and in magnitude is about three-fourths the decadal decline found through most of the adult age range (Clifton et al. 2016: Table 1 and Figure 1A).
4. <u>Consistency across waves</u>: No opportunity to test.
5. <u>Sexuality</u>: No opportunity to test.
6. <u>Diabetes</u>: An age-adjusted association of T with self-reported diabetes for men approached significance (p = .09) but this did not persist after adjusting for other confounding factors including BMI.
7. <u>Morbidity</u>: No opportunity to test mortality against T. Salivary T was associated, independently of age, with a range of measures of general health in men. Cardiovascular disease (including hypertension) was significantly associated with lower mean T in men, independently of age. There was weak evidence of an association between lower T and having a long-standing illness or disability (p=0.07) and strong evidence of an association between lower salivary T and having 2 or more comorbid health conditions (p=0.01), though no significant association between mean salivary T and self-reported general health.

Overall, correlations what could be tested with the British Natsal-3 data, in which salivary T was assayed by LC-MS/MS, were replicated. An exception was the previously reported association of T with type 2 diabetes and HbA1c, which was not replicated in either dataset.

## DISCUSSION

Our original intent was to see if several relationships involving male T, previously reported in the literature for a range of adult ages, held true among older men whose T was assayed from saliva. To that end, we tested these relationships on older men in two datasets, the American NSHAP, where salivary T was measured by EIA, and the British Natsal-3, where salivary T was measured by LC-MS/MS.

The expected correlations did not, or only weakly, replicated with the American data but did replicate with the British data. The lack of a diurnal decline in T among NSHAP’s American men was especially surprising because it has been reported so often. Timing of saliva sampling was dictated by the times of interviews, which varied throughout the day. Possibly there were extraneous but unknown differences between men interviewed in the morning and those interviewed later in the day.

The British Natsal-3 data provide strong evidence that previously reported correlations, at least those we could test, continue to hold among older men, and that saliva is an acceptable medium for assaying T. This presumes that the assay technique, in this case liquid chromatography-tandem mass spectroscopy (LC-MS/MS), is accurate. Why, then, did the correlations fail to replicate, or at best only weakly, in the American NSHAP data? One possibility is that there was considerable noise in the American values for T, which obscured correlations that might in fact exist.

Although EIA of salivary T is today widely used in biobehavioral studies, we must reconsider its accuracy for measuring salivary T. Recently Welker, et al. (2016) found measures of salivary T differed considerably across EIAs of three commonly used kit manufacturers. They also found notable differences between EIA values and those assayed by LC-MS/MS, a technique noted for its specificity. EIA tends to inflate estimates of lower T concentrations in women; and EIA values for salivary T, while modestly correlated with serum levels for males, are less so for females (Shirtcliff, Granger and Likos 2002).

In women, circulating concentrations of T are typically around 5–10% of those in men (Haring et al. 2012; Davison et al. 2005). However, EIA often gives sex ratios far lower. For example, the three EIA manufacturers used by Welker, et al. (2016, supplementary table S3) gave sex ratios for salivary T at 2.4, 2.3, and 2.0, whereas the sex ratio given by their LC-MS/MS was 5.5.

Turning to the data used in this paper, in NSHAP, the male-to-female ratio as measured by EIA = 1.8 in both waves. In the Natsal-3 survey of Britons, the male-to-female ratio as measured by LC-MS/MS is about 6 for the full age range and nearly the same for respondents age 57+ (Keevil, et al. 2017). Comparing T in NSHAP with T in Natal-3 respondents age 57+ suggests that for men, EIA estimated mean T nearly two times higher than LC-MS/MS did. For women, EIA estimated mean T nearly six times higher than LC-MS/MS did.

Calculations from supplementary table S3 of Welker, et al. (2016) show the same pattern, though less extreme. That is, using LC-MS/MS as a standard, EIA estimates of mean T are over twice as high for women as for men. These results suggest that EIA has unrecognized cross-reactivity that may swamp the low T concentrations typical of females. Any such crossreactivity would be less problematic for men with their higher T levels, but it may introduce sufficient error to obscure real correlations, disguising them as nulls.

The major shortcoming of our criticism of EIA is that the two assay methods were used on different samples. Obviously, a stronger test would have used both assays on the same sample, but we had to use such surveys as were available. We believe that corroborating important correlations in a sample using LC-MS/MS, but not finding them in a sample using EIA, importantly implicates the inaccuracy of EIA for salivary testosterone.

In 2013, the *Journal of Clinical Endocrinology & Metabolism* considered a requirement for mass spectrometry sex steroid assays (Handelsman and Wartofsky 2013), although an explicit ban on enzyme immunoassay of sex steroids is not presently in place. Nonetheless, this indicated the disquiet among researchers about EIA measurements of sex steroids and their unreliable findings.

The existing literature on social neuroendocrinology in humans is rife with inconsistent findings regarding testosterone (Mazur 2018). In recent decades these reports have depended increasingly on EIA. In view of evidence for the inadequacy of these assays, it is time for a review of past work, flagging results dependent on EIA for possible distortions they may have introduced into the literature. We may need to go back to square one.

